# GAVISUNK: Genome assembly validation via inter-SUNK distances in Oxford Nanopore reads

**DOI:** 10.1101/2022.06.17.496619

**Authors:** Philip C. Dishuck, Allison N. Rozanski, Glennis A. Logsdon, Evan E. Eichler

**Affiliations:** Department of Genome Sciences, University of Washington School of Medicine, Seattle, WA 98195, USA; Howard Hughes Medical Institute, University of Washington, Seattle, WA 98195, USA

## Abstract

**Motivation:** Highly contiguous *de novo* genome assemblies are now feasible for large numbers of species and individuals. Methods are needed to validate assembly accuracy and detect misassemblies with orthologous sequencing data to allow for confident downstream analyses.

**Results:** We developed GAVISUNK, an open-source pipeline that detects misassemblies and produces a set of reliable regions genome-wide by assessing concordance of distances between unique *k-*mers in Pacific Biosciences high-fidelity (HiFi) assemblies and raw Oxford Nanopore Technologies reads.

**Availability:** GAVISUNK is available at https://github.com/pdishuck/GAVISUNK.

## 1 Introduction

Highly accurate and contiguous *de novo* assemblies of long-read sequencing data have made reference-grade assemblies feasible for many species and individuals (Rhie *et al*., 2021; Ebert *et al*., 2021). Pacific Biosciences high-fidelity (HiFi) sequencing in particular has facilitated some of the first complete assembly of centromeres, acrocentric regions, as well as other complex segmental duplications (Miga *et al*., 2020; Logsdon *et al*., 2021; Nurk *et al*., 2022). These assemblies make possible comprehensive, whole-genome evaluations of sequence variation, including some of the most difficult-to-assemble regions, unbiased by reference alignments for the first time. Phased genome assemblies, however, are still subject to the collapse of repetitive sequences, incorrect orientations, and misassemblies. Thus, any discoveries based on these automated shotgun sequence assemblies raise the question of whether the assembled sequence is, in fact, valid.

Here, we present genome assembly validation via inter-singly unique *k*-mer (SUNK) distances in Oxford Nanopore Technologies (ONT) reads, known hereafter as GAVISUNK. GAVISUNK is a method of validating HiFi-driven assemblies with orthogonal ONT sequence. It specifically assesses the contiguity of regions, flagging potential haplotype switches or misassemblies. Although the ONT platform has a significantly higher error rate than that of HiFi (Logsdon *et al*., 2020), such reads are typically much longer, making it a powerful orthogonal approach for assessing both contiguity and read depth across regions of interest.

Whereas previous genome blacklists or masks of inaccessible regions, such as those used by the ENCODE Consortium, are determined based on annotation of a reference genome (Amemiya *et al*., 2019), GAVISUNK may be applied to any region or genome assembly to identify misassemblies and potential collapses and is, thus, particularly valuable for validating the integrity of regions with large and highly identical repeats that are more prone to assembly error. This method can be applied genome-wide or at fine scale to closely examine regions of interest across multiple haplotype assemblies.

## 2 Methods

This assembly validation method relies on identifying SUNKs, *k*-mers that occur just a single time within the HiFi-based assembly (Sudmant *et al*., 2010) and confirming these SUNKs within long ONT sequencing reads. Because of the relatively lower accuracy of ONT data, false SUNK overlaps may occur at an appreciable frequency between paralogous regions of the genome or haplotypes. Therefore, we leverage not only the presence or absence of SUNKs, but the intervening distance between pairs of SUNKs, referred to hereafter as inter-SUNK distances. The approach thus compares the expected inter-SUNK distance in the assembly and observed distance within ONT reads to recruit reads to their corresponding genomic location. The failure of ONT reads to span between SUNKs in the assembly defines a misjoin, while an excess of reads flags a potential collapse.

We apply Jellyfish (Marçais and Kingsford, 2011) to identify unique *k*-mers based on all HiFi contigs within an assembly, generating a set of SUNKs for validation. ONT reads are haplotype-phased using parental Illumina whole-genome sequencing data via Canu (*Koren et al*., 2017). The position of all SUNKs within each ONT read are identified and used for downstream analysis. Each ONT read is assigned to its best-matching HiFi assembled contig and orientation by comparing the locations of read SUNKs to assembly SUNKs within a diagonal band centered on the median SUNK location for that read. SUNKs observed in ONT reads at an implausibly high or low frequency (i.e., greater than four standard deviations above the mean or fewer than twice) for either haplotype are excluded from consideration.

To validate the assembly of each HiFi contig, all reads identified in the previous step are considered. For each read, a matrix of all pairwise inter-SUNK distances within the read is generated using NumPy and compared to expected distances from the assembly, allowing +/-2% variation in length for a given distance by default (Harris *et al*., 2020). Only the largest set of SUNKs with fully inter-consistent inter-SUNK distances are retained from each read for subsequent validation. In this way, chimeric reads and other spuriously connected SUNKs are separated. A graph is generated for each contig, with SUNKs as nodes and reads as edges, connecting pairs of SUNKs with consistent inter-SUNK distances, using the graph-tool library. This graph is decomposed into its connected components, each of which now corresponds to a validated region of the contig, identifying SUNKs spanned by ONT reads. These validated regions are sent as output to a BED file, and assembly SUNKs with no read support are listed as potentially artifactual.

## 3 Usage and Examples

GAVISUNK can run on consumer hardware, research clusters, or in cloud computing environments, as its steps are automated as part of a configurable Snakemake workflow (Mölder *et al*., 2021). To install GAVISUNK, clone the repository at https://github.com/pdishuck/GAVISUNK. Upon first run, dependencies will install automatically as a conda environment. As described in the README, two configuration files must be edited to specify the assembly and *k-*mer size (`config.yaml`) and input nanopore read files (`ont.tsv`). To execute the pipeline, submit

`snakemake –use-conda –cores {thread.count}`

By default, the pipeline produces a BED file of validated regions (`hap#.validated.bed`) and the complementary unconfirmed “gaps” between validated regions (`hap#.gaps.bed`) for each haplotype of the assembly. Summary outputs from example assemblies are shown in **Table 1**. For the region of each validation gap, the pipeline produces a visualization to show SUNK-tagged read support, in PDF, SVG, and PNG formats (`contig_start_end.format`), as shown in **Figure 1**. Optionally, a BED file with regions of interest for visualization can be specified for each haplotype in `ont.tsv`.

**Table 1.**
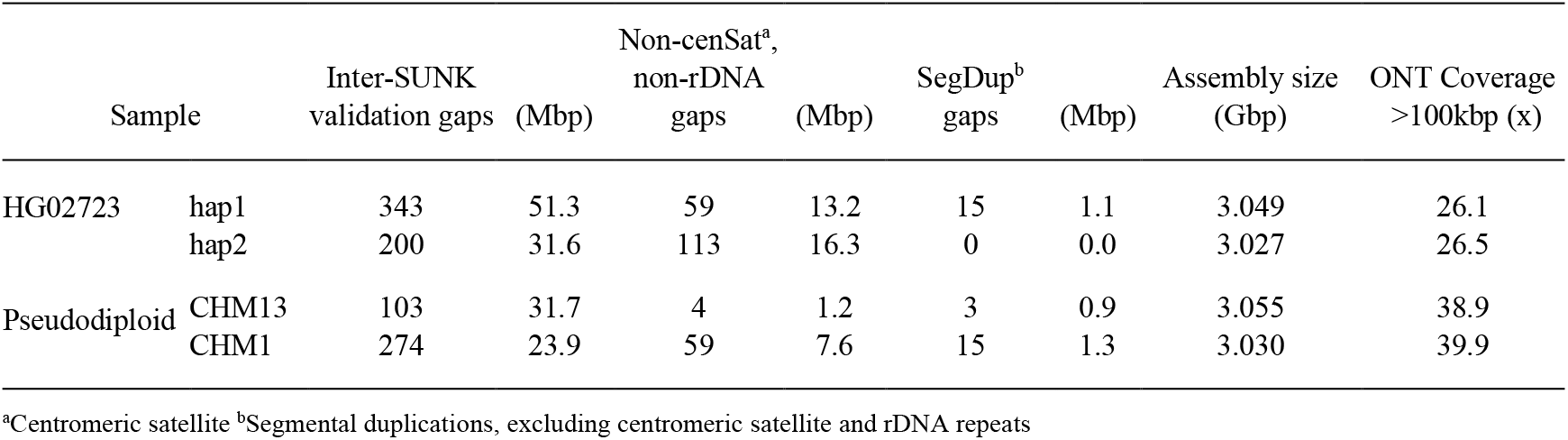
Results of genome-wide analysis.

**Figure 1.**
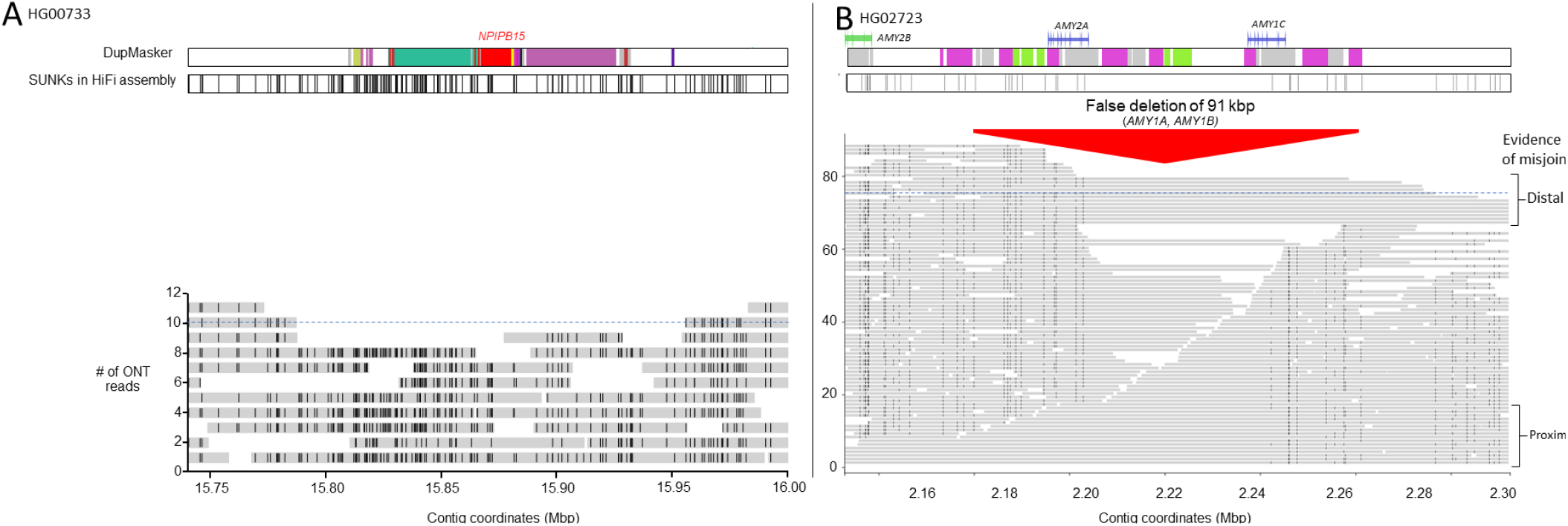
**A**. An example of a validated region within the maternal haplotype assembly of HG00733. Horizontal gray bars represent individual ONT reads, and vertical black lines indicate the position of SUNKs originally detected from the HiFi assembly. All possible assembly SUNKs are shown above, along with a DupMasker track marking segmental duplications. The dotted blue line indicates the mean genome-wide coverage of ONT sequencing data. **B**. An example of a misassembly within the paternal haplotype assembly of HG02723 detected by GAVISUNK. This misassembly resides within the amylase (*AMY*) duplication locus and contains a false deletion of *AMY1A* and *AMY1B*, which is confirmed by the failure of ONT to anchor across the misjoin.

To annotate the visualizations, BED files encoding regions of interest and color may be supplied. In the example shown in **Figure 1**, DupMasker annotations demarcate segmental duplications across the locus (Jiang *et al*., 2008). Additionally, the sizes of unspanned inter-SUNK gaps are output for comparison with the distribution of inter-SUNK distances and expectation of spanning that distance given empirical ONT coverage at that read length (**Supplementary Figures 1**,**2**).

We constructed a pseudo-diploid genome assembly as a benchmark of false discovery rate for a well-curated reference assembly with matched ultra-long ONT data. We used the long-read assemblies of the CHM13 and CHM1 human cell lines, both of which are derived from complete hydatidiform moles exhibiting genome-wide uniparental disomy and are therefore guaranteed to represent a single haplotype. CHM13 was assembled by the T2T Consortium with a variety of methods and manual validation (Nurk *et al*., 2022), and CHM1 was assembled from HiFi data (Vollger *et al*., 2022 using hifiasm v0.12 (Cheng *et al*., 2021)), each with >30x coverage of ONT reads longer than 100 kbp. Results for this pseudo-diploid, along with true diploid HG02723 assembled with hifiasm v0.14 (GenBank GCA_018504065.1), are summarized in **Table 1**. Long repetitive arrays of centromeric satellite sequence and rDNA often fail to validate with this method, but ongoing work aims to optimize GAVISUNK for usage in these regions. For CHM13, 1.0% of the assembled genome (103 gaps, 31.7 Mbp) is unsupported by ONT inter-SUNK distances, and 0.7% (274 gaps, 23.9 Mbp) for CHM1. For comparison, assuming random distribution of the empirical ONT read lengths compared to the distances between SUNKs for the CHM13 assembly, 24.9 Mbp is expected to be unsupported, corresponding to 92 gaps.

## 4 Conclusion

We developed GAVISUNK, a method for assembly validation using inter-SUNK distances in ONT reads. Applied to HiFi genome assemblies, this tool provides orthogonal validation of regions for downstream analysis, allowing for subsequent genome analyses and annotation.

GAVISUNK provides easily interoperable BED outputs and interpretable visualizations of supported and unsupported regions of interest.

## Acknowledgements

We thank Mitchell Vollger and William Harvey for their helpful algorithmic, programming, and source control advice, and Tonia Brown for proofreading. This article is subject to HHMI’s Open Access to Publications policy. HHMI lab heads have previously granted a nonexclusive CC BY 4.0 license to the public and a sublicensable license to HHMI in their research articles. Pursuant to those licenses, the author-accepted manuscript of this article can be made freely available under a CC BY 4.0 license immediately upon publication.

---

**Supplementary Figure 1.**
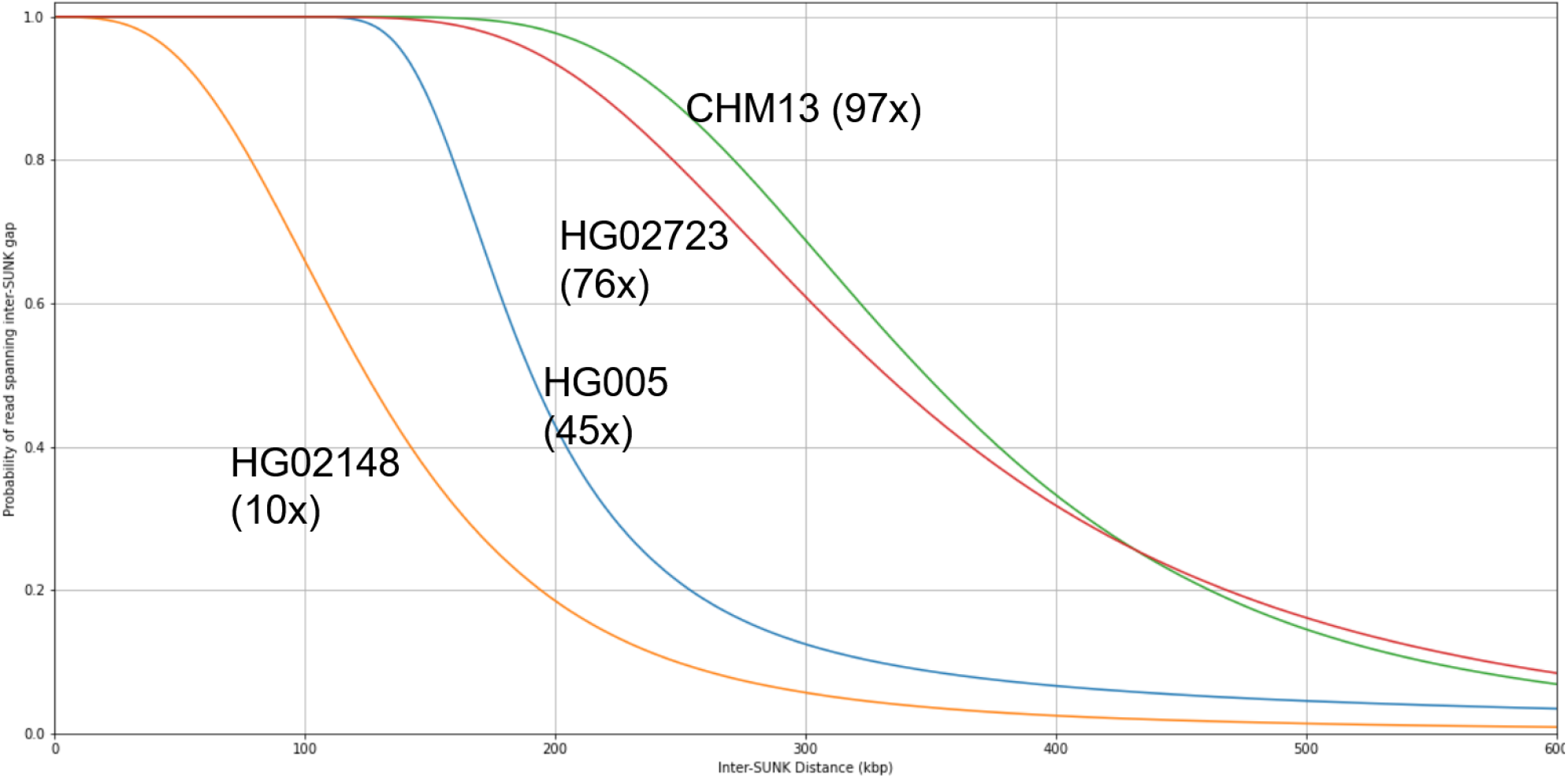
Simulation of probability of spanning inter-SUNK gaps. The probability of reads spanning a given inter-SUNK distance is determined on a per-sample basis by calculating the Poisson distribution of ONT reads across the genome, adjusting for sequencing accuracy and size of the SUNK group. Haplotype-specific ONT sequencing depth is shown in parentheses for each sample.

**Supplementary Figure 2.**
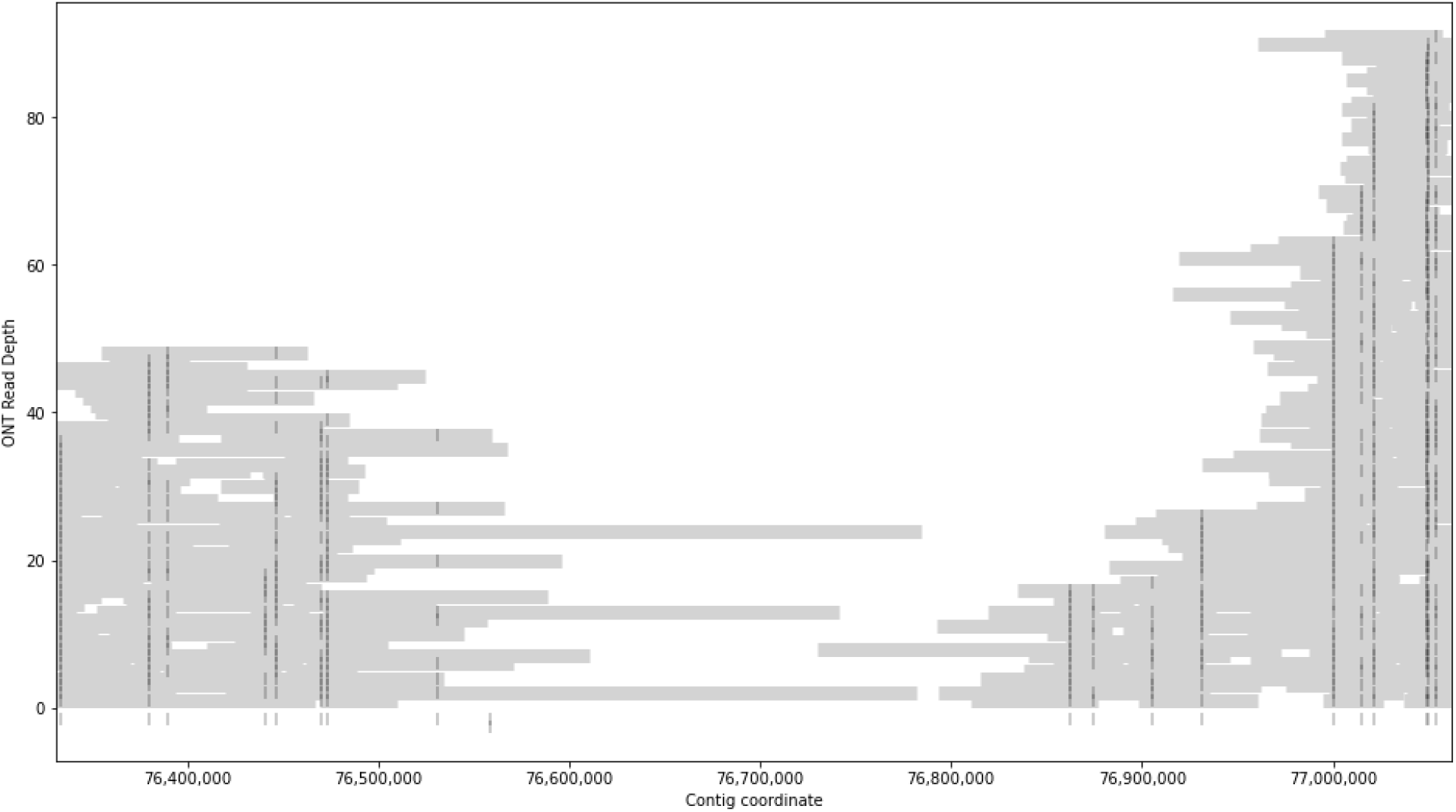
Unspannable gap in CHM13-T2T *HYDIN*. Because the SUNK density at this recently duplicated locus is too low for ONT reads at the given length and coverage to span between SUNK groups, no determination can be made of assembly correctness.

